# Marker effect p-values for single-step GWAS with the algorithm for proven and young in large genotyped populations

**DOI:** 10.1101/2023.10.15.562399

**Authors:** Natália Galoro Leite, Matias Bermann, Shogo Tsuruta, Ignacy Misztal, Daniela Lourenco

## Abstract

**Background:** Although single-step GBLUP (**ssGBLUP**) is a breeding value method, single-nucleotide polymorphism (**SNP**) effects can be backsolved from ssGBLUP genomic estimated breeding values (GEBV), and p-values can be obtained as a measure of estimation certainty. This enables single-step genome-wide association studies (**ssGWAS**). However, obtaining p-values for ssGWAS relies on the inversion of dense matrices, which poses computational limitations in large genotyped populations. In this study, we present an algorithm to approximate p-values for SNP in ssGWAS with many genotyped animals. The approximation relies on the algorithm for proven and young (APY) and submatrices for core animals. To test that, we first compared SNP p-values obtained with an exact inversion using the genomic relationship matrix (**G**^**−1**^) for 50K genotyped animals to those estimated with an exact inversion using 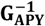 and those obtained with the proposed approximation based on 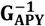. Then, we compared these results with those obtained with the proposed approximation using 450K genotyped animals.

**Results:** The same genomic regions in chromosomes 7 and 20 were identified with p-values obtained with **G**^**−1**^, 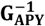, and the approximation based on 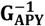 when using 50k genotyped animals and 1.5M in the pedigree. In terms of computational requirements, obtaining p-values with the approximation based on 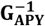 represented a reduction of 38 times in wall-clock time and ten times in memory requirement compared to using the exact inversion with 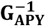. When the approximation was applied to a population of 450K genotyped animals and 1.8 in the pedigree, apart from the two genomic regions in chromosomes 7 and 20 previously identified with the smaller genotyped population, two new significant regions on chromosomes 6 and 14 were uncovered, indicating an increase in GWAS detection power when including more genotypes in the analyses. The process of obtaining p-value with the approximation and 450K genotyped individuals took 24.5 wall-clock hours and 87.66 GB of memory, which is expected to increase linearly with the addition of noncore genotyped individuals.

**Conclusions:** With an algorithm that approximates the prediction error variance of SNP effects based on APY, ssGWAS with p-values for SNP is possible in large genotyped populations. The computational cost of obtaining p-values in ssGWAS is no longer a limitation in extensive populations with many genotyped animals.

## Background

The single-step genomic best linear unbiased prediction (**ssGBLUP**) has been successfully implemented in the routine genetic evaluation of several livestock species (Lourenco et al., 2015; Tsuruta et al., 2021; Abdollahi-Arpanahi et al., 2022). The vast adoption of ssGBLUP is associated with the straightforward, simultaneous evaluation of populations composed of genotyped and non-genotyped animals, the non-requirement of pseudo phenotypes, the decrease in biases attributed to double counting and genomic preselection, and reliable estimation of breeding values for complex genetic models (Misztal et al., 2020).

Although ssGBLUP is a breeding value-based method that provides genomic EBV (**GEBV**), obtaining single-nucleotide polymorphism (**SNP**) effects from this method may also be valuable when investigating how genome segments are associated with important traits. When that is the case, SNP effects can be easily obtained from a linear transformation of GEBV following formulas presented by VanRaden (2008), Strandén and Garrick (2009), and Wang et al. (2012). Besides SNP effects, the proportion of genetic variance explained by single SNP or by SNP windows can help identify important regions of the genome in single-step genome-wide association studies (**ssGWAS**) (Fragomeni et al., 2014; Misztal et al., 2014c; Wang et al., 2014). However, this procedure does not consider the uncertainty of the SNP effect estimation, making it more difficult to replicate findings from ssGWAS (Fragomeni et al., 2014; Wang et al., 2014). To overcome this problem, Aguilar et al. (2019) presented formulas for obtaining frequentist p-values for ssGWAS as an extension of the ideas previously presented by Gualdrón Duarte et al. (2014), Bernal Rubio et al. (2016), and Lu et al. (2018) in the ssGBLUP context. The authors also showed that p-values could be successfully obtained within a reasonable computational time for a large Angus population accounting for roughly 1M phenotyped individuals, 1,500 genotyped sires, and about 1.8M animals in the pedigree.

The formulas presented by Aguilar et al. (2019) require obtaining the prediction error variance of the SNP effects (**PEV**_**SNP**_) which relies on obtaining the breeding value prediction error (co)variance for genotyped animals (**C**^**u2u2**^) through the inversion of the left-hand side of the mixed model equations (**LHS**). The inversion of such a matrix becomes challenging as the number of traits and animals increases. With genomic information, the LHS contains a very dense block represented by the inverse of the genomic relationship matrix (**G**^**−1**^), which is hard to obtain directly for more than 100K genotyped animals (Misztal et al., 2020). One approach to deal with the computation limits with large genotyped populations is to use a sparse representation of **G**^**−1**^ created by the algorithm for proven and young (**APY**) (Misztal et al., 2014a). In APY, the genotyped individuals are split into two sets. The set of genotyped animals representing all genomic variation is called ‘core’ (non-redundant information); the remaining animals are ‘noncore’ (redundant information). Then, the GEBV of noncore animals are conditioned on the GEBV of core animals, making **G**^**−1**^ very sparse. Apart from increasing the sparseness of **G**^**−1**^, Bermann et al. (2022a) showed that, with APY, obtaining PEV_SNP_ can be reduced to components only associated with the PEV of GEBV for the core set (**PEV**_**CC**_), drastically reducing the dimensionality of matrices involved in calculations to obtain PEV_SNP_. However, in the formulas shown by Bermann et al. (2022a), obtaining PEV_CC_ still requires a direct inversion of the LHS with all genotyped animals. Even though **G**^**−1**^ is sparser with APY, components in the LHS of single-step equations such as the inverse of the pedigree relationship matrix for genotyped animals 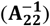 are still dense, thus implying that computation limits for the inversion of LHS might still exist.

One way to overcome this problem is to obtain an approximated prediction error variance of breeding values for the APY core set (**PEV**_**CCappx**_) that does not require the inversion of the LHS. For that, an extension of the algorithm proposed by Misztal and Wiggans (1988) that accounts for genomic information with APY was presented by Bermann et al. (2022b). In this algorithm, Bermann et al. (2022b) showed that PEV_CCappx_ can be obtained with a block-sparse inversion of **G**^**−1**^ with APY 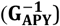 plus a diagonal matrix of contributions from phenotypes and pedigree relationships. Empirical results shown by the authors demonstrate that PEV_CCappx_ is obtained in a few minutes for an Angus population with about 300K genotyped animals. Moreover, they showed that, although computation complexity increases cubically with the number of core animals, that remains linear for the noncore set. Thus, by combining APY and PEV_CCappx_, it may be possible to approximate p-values for SNP and enable ssGWAS for large genotyped populations while lifting the current computational limitations. Therefore, this study presents an algorithm to approximate p-values for SNP in ssGWAS with many genotyped animals based on APY. The performance of the proposed algorithm was tested against the regular way to compute p-values using the exact inverse of LHS with **G**^**−1**^ or 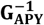 with 50K genotyped animals. Then, the final test involved applying the proposed algorithm to a dataset with 450K genotyped animals.

## Methods

### Theory

In ssGBLUP, SNP effects can be obtained from backsolving GEBV using a linear transformation (VanRaden, 2008; Strandén and Garrick, 2009; Wang et al., 2012):

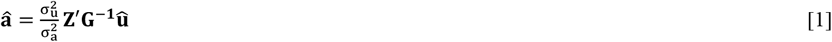

where **â** is the vector of SNP effects, 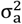 is SNP variance, 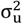 is the genetic variance, **û** is the vector of breeding values, **Z** is a matrix of SNP content centered by two times the allele frequency (p), and **G**^**−**1^ is the inverse of the genomic relationship matrix, with **G** constructed as the type I of VanRaden (2008):

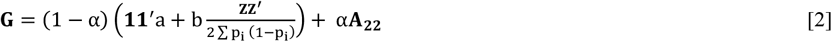

where α is the blending parameter (5%) to avoid singularity problems in **G** (VanRaden, 2008); a and b are tuning parameters to assure the compatibility between **G** and **A**_22_ (Vitezica et al., 2011), and other elements were defined above.

Once SNP effects (**â**) are estimated, the p-value for the ith SNP can be obtained as shown by Aguilar et al. (2019):

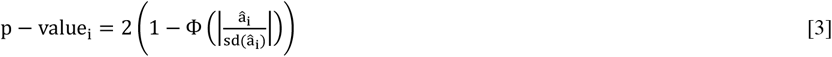

where sd(â_i_) is the square root of the variance of the ith SNP effect estimate obtained as (Gualdrón Duarte et al., 2014):

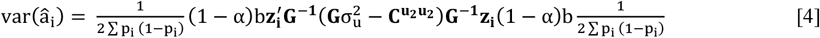

with 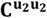 referring to the matrix of breeding value prediction error (co)variance for genotyped animals, and other parameters are defined above. The computation of var(â_i_) is restrained by the costs associated with obtaining **G**^**−1**^ and **C**^**u2u2**^. Those components result from the inversion of dense matrices of high dimension, and obtaining them becomes unfeasible with large genotyped populations (Aguilar et al., 2019; Garcia et al., 2020).

The computational limitations of obtaining **G**^**−1**^ can be overcome by replacing this matrix with its sparse representation built with **APY**(Misztal et al., 2014a). With APY, a small set of genotyped animals (core) is chosen, and the relationship of the remaining animals (noncore) is obtained by recursive equations on the core set with linear computing cost. The inverse of the genomic relationship matrix with APY is constructed as follows:

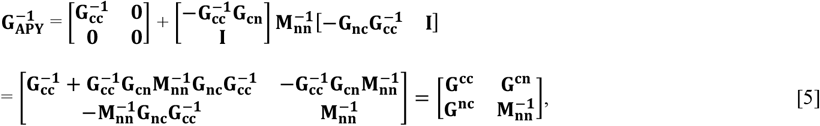

where 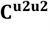 and 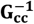 are the inverses of the full genomic relationship matrix for core and diagonal for noncore animals, respectively, and **G**_**cn**_ is the genomic relationship matrix between core and noncore animals. The elements of the matrix 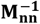 are obtained as:

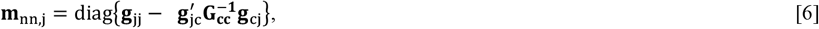

where g_jj_ is the diagonal element of **G**_**nn**_ for the jth animal, and **g**_jc_ is the relationship between the jth noncore animal with core animals. With APY, the need to invert a dense and high dimensional **G** is reduced to only inverting the genomic relationship matrix for core animals (**G**_**cc**_) [Eq. 5], which for most livestock species or breeds contains less than 15K animals (Pocrnic et al., 2016; Cesarani et al., 2022).

Beyond the reductions in computing costs of obtaining **G**^**−1**^, Bermann et al. (2022a) showed that, with APY, estimating the PEV_SNP_ is reduced to components only associated with the core animals:

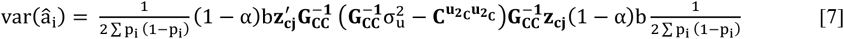

where **z**_**cj**_ is the jth line of the **Z** matrix for core animals, and 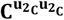 is the prediction error covariance matrix of breeding values for core animals, and other elements are as defined before. However, obtaining 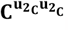, which we can also refer to as **PEV**_**CC**_, still depends on the inversion of a high dimension LHS, which might yet limit computations as model complexity and the number of genotyped animals increase. To overcome this limitation, an approximation of the PEV_CC_ (**PEV**_**CCappx**_) can be obtained as follows (Bermann et al., 2022b):

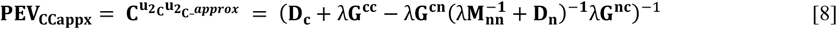

where 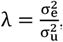, 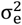 is the residual variance, and **D** and **D** are the blocks for core and noncore animals from the diagonal matrix **D** constructed as (VanRaden and Freeman, 1985; Misztal and Wiggans, 1988):

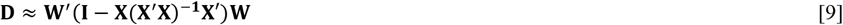

where **W** and **X** are incidence matrices for animal and fixed effects. Therefore, var(â_i_) when combining APY and PEV_CCappx_ is:

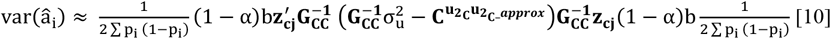

where all parameters were defined above. Equation [10] implies that when APY and PEV_CCappx_ are combined, SNP p-values can be calculated with matrices only associated with core animals and without the requirement of inverting the LHS, thus potentially lifting the current computational limitations for large genotyped populations.

Therefore, the proposed algorithm to approximate p-values for SNP involves the following steps:

1. Save 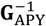 and components of 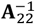 in disk (PREGSF90 from BLUPF90 software suite; Misztal et al. (2014b));
2. Obtain GEBV based on APY, reading the saved matrices (BLUP90IOD3 from BLUPF90 software suite);
3. Compute PEV_CCappx_ using block sparse inversion as in Bermann et al. (2022b) (ACCF90GS2 from BLUPF90 software suite);
4. Use PEV_CCappx_ and 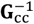 from 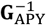 to compute var(â_i_) as in Equation [10] (POSTGSF90 from BLUPF90 software suite);
5. Compute SNP p-values_i_ as in Equation [3] by using the square root of var(â_i_) obtained in step 4 (POSTGSF90 from BLUPF90 software suite).

### Dataset

The American Angus Association (Saint Joseph, MO) provided the dataset to test the proposed algorithm to approximate p-values for SNP. A total of 844,726 animals born from 2012 to 2017 were scored for post-weaning gain (**PWG**). Phenotyped animals were produced by 93,161 sires and 812,292 cows and were distributed into 64,889 contemporary groups. Genomic information on 39,744 SNP (after quality control) was available for 450,673 animals born from 2012 to 2018. Of the genotyped animals, 217,434 were also phenotyped, whereas the remaining animals only contributed with genotypes and pedigree. Pedigree information was available for all phenotyped and genotyped animals up to 3 generations of relationships, summing up 1,837,789 records.

### Statistical model

A single-trait animal model was used for the estimation of PWG breeding values as follows:

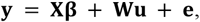

where **y** is the vector of PWG phenotypes; **β** is the vector containing the fixed effect of contemporary groups; **u** is the vector of random additive genetic effects; **e** is the vector of random residuals; and **X**, and **W** are incidence matrices for the effects contained in **β** and **u**, respectively. Random effects were distributed as 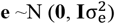 and 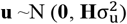, where **I** is an identity matrix, and **H** is the realized relationship matrix for genotyped and non-genotyped animals in ssGBLUP, with inverse constructed as shown by Aguilar et al. (2010):

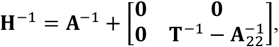

where **A**^**−**1^ is the inverse of the pedigree relationship matrix and **T**^**−**1^ is equal to 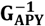 [Eq. 5] for genetic analyses with APY, and equal to **G**^**−1**^ [Eq. 2] otherwise. The 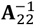 was defined before.

### Statistical analyses

#### Comparison between p-value computing methods with a reduced dataset

In this set of analyses, we aimed to compare exact p-values obtained with a regular **G**^**−1**^ (**Exact_Ginv**) as a benchmark [Eq. 3, 4], with 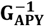 and exact PEV_cc_ (**Exact_GinvAPY**) [Eq. 3, 7], and p-values with 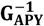 and PEV_CCappx_ (**Approx_GinvAPY**) [Eq. 3, 10]. Because the p-values from Exact_Ginv and Exact_GinvAPY require obtaining the inverses of the genomic relationship matrix and of the LHS, a reduced subset of 50K randomly selected genotyped animals among the 450K was used for all p-value computing methods to ensure computation feasibility and fair comparisons. As the 50K genotyped animals were selected randomly, sampling was repeated three times. Phenotypic information was kept complete for all p-value computing methods, but the number of animals in the pedigree slightly varied across samples.

The method comparison was based on the Pearson correlation of computed p-values, SNP effects, var(â_i_), in addition to visual inspection of Manhattan plots. Wall-clock time and memory requirements were also recorded. Analyses involving APY were composed of a core set of 13,030 genotyped animals, which corresponded to the number of eigenvalues explaining 98% of the genetic variance in **G** with all genotypes available (i.e., 450,673 genotyped animals) (Pocrnic et al., 2016).

#### Application of Approx_GinvAPY in a large genotype population

In the second set of analyses, we aimed to calculate p-values with 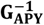 and PEV_CCappx_ with the full set of 450K genotyped animals (**Approx_GinvAPY450K**). For straightforward interpretations, the core sets in Approx_GinvAPY450K were composed of the same set of animals as in the analyses with 50K genotypes. For example, within the same replicate, the core set in Exact_Ginv, Approx_GinvAPY, and Approx_GinvAPY450K comprised the same genotyped animals. Elapsed wall-clock time and memory requirements were recorded, and visual inspection of Manhattan plots helped to evaluate the soundness of the approximation. A summary of the information available for all analyses is in Table 1.

**Table 1.** Number of records per source of information for all scenarios.

| Method<sup>a</sup> | Genotypes | Core | Pedigree<sup>b</sup> | Phenotypes |
| --- | --- | --- | --- | --- |
| Exact_Ginv | 50,000 | 13,030 | 1,576,738 | 844,726 |
| Exact_GinvAPY | 50,000 | 13,030 | 1,576,738 | 844,726 |
| Approx_GinvAPY | 50,000 | 13,030 | 1,576,738 | 844,726 |
| Approx_GinvAPY450K | 450,673 | 13,030 | 1,837,789 | 844,726 |

<sup>a</sup>SNP p-values obtained from a data set of 50K genotyped animals with an exact inversion of the genomic relationship matrix ( $\mathbf{G}^{-1}$ ; (Exact\_Ginv), with a sparse $\mathbf{G}^{-1}$ build with the algorithm for proven and young ( $\mathbf{G}_{APY}^{-1}$ ; Exact\_GinvAPY), and from an approximation based on $\mathbf{G}_{APY}^{-1}$ (Approx\_GinvAPY). Approx\_GinvAPY450K refers to the Approx\_GinvAPY method applied to a genotyped population of 450K animals.
<sup>b</sup>Because pedigree traced back three generations of relationships from phenotyped and genotyped animals, the number of animals in the pedigree slightly varied from 1,576,112 to 1,837,789 among replicates and scenarios.

All analyses were performed with software from the BLUPF90 software suite (Misztal et al., 2014b) on an Intel(R) Xeon(R) CPU E5-2650 v4 @ 2.20GHz server with 24 threads. New implementations for obtaining p-values with 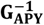 and PEV_CCappx_ were available in modified versions of BLUP90IOD3 and ACCF90GS2 (Lourenco et al., 2022).

## Results and Discussion

Using ssGWAS for association studies in farm animal populations increases the detection power because it considers phenotypic information from non-genotyped individuals, allows for complex models involving multiple traits and environmental and genetic correlated effects, and does not rely on pseudo phenotypes (Legarra et al., 2014; Wang et al., 2014; Mancin et al., 2021). For ssGWAS, SNP effects from ssGBLUP are easily obtained by backsolving GEBV already routinely estimated in the breeding programs (Strandén and Garrick, 2009; Wang et al., 2012). Similarly, the prediction error (co)variance of GEBV is backsolved to the variance of estimating SNP effects, which is used to compute SNP p-values (Aguilar et al., 2019). Before the availability of SNP p-values in ssGWAS, the important genomic regions were associated with single SNPs or SNP windows that explained more than 1% of the genetic variance (Fragomeni et al., 2014; Garcia et al., 2018). However, this approach relies on the frequency of SNPs and assumes that SNP effects are estimated without error, which may challenge the persistence of results over time and among different statistical methods (Fragomeni et al., 2014; Misztal et al., 2014c). Therefore, having p-values for ssGWAS is of great importance for association studies. Nonetheless, the method to compute SNP p-values presented by depends on the inversion of the LHS and should become prohibited with increasing data dimensionality. In this study, we approach the challenge of computing p-value in populations with an increasing number of genotyped individuals. For that, we compared three methods to calculate p-values, which consisted of obtaining p-values with (1) a regular **G**^**−1**^ (Exact_Ginv), (2) with 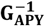 and exact PEV_cc_ (Exact_GinvAPY), and (3) an efficient algorithm combining 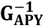 and PEV_CCappx_ (Approx_GinvAPY). We later evaluated the performance of the Approx_GinvAPY method when applied to a large genotyped population comprised of around 450K individuals (Approx_GinvAPY450K).

### Comparison between p-values computing methods with a reduced dataset

Manhattan plots for all investigated p-values computing methods are shown in Figures 1-3. Across all methods and replicates, two significant peaks were identified in chromosomes 7 and 20 for PWG. The top three SNP in each peak were overall correctly identified for all APY computing methods with respect to Exact_Ginv (Figures 1-3). The correct identification of the same genome regions with all p-values computing methods indicates that (1) 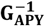 is a proper sparse representation of **G**^**−1**^, and (2) combining 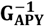 and PEV_CCappx_ results in accurate results in comparison with Exact_Ginv, making APY suitable for ssGWAS analyses for large genotyped populations.

**Figure 1.**
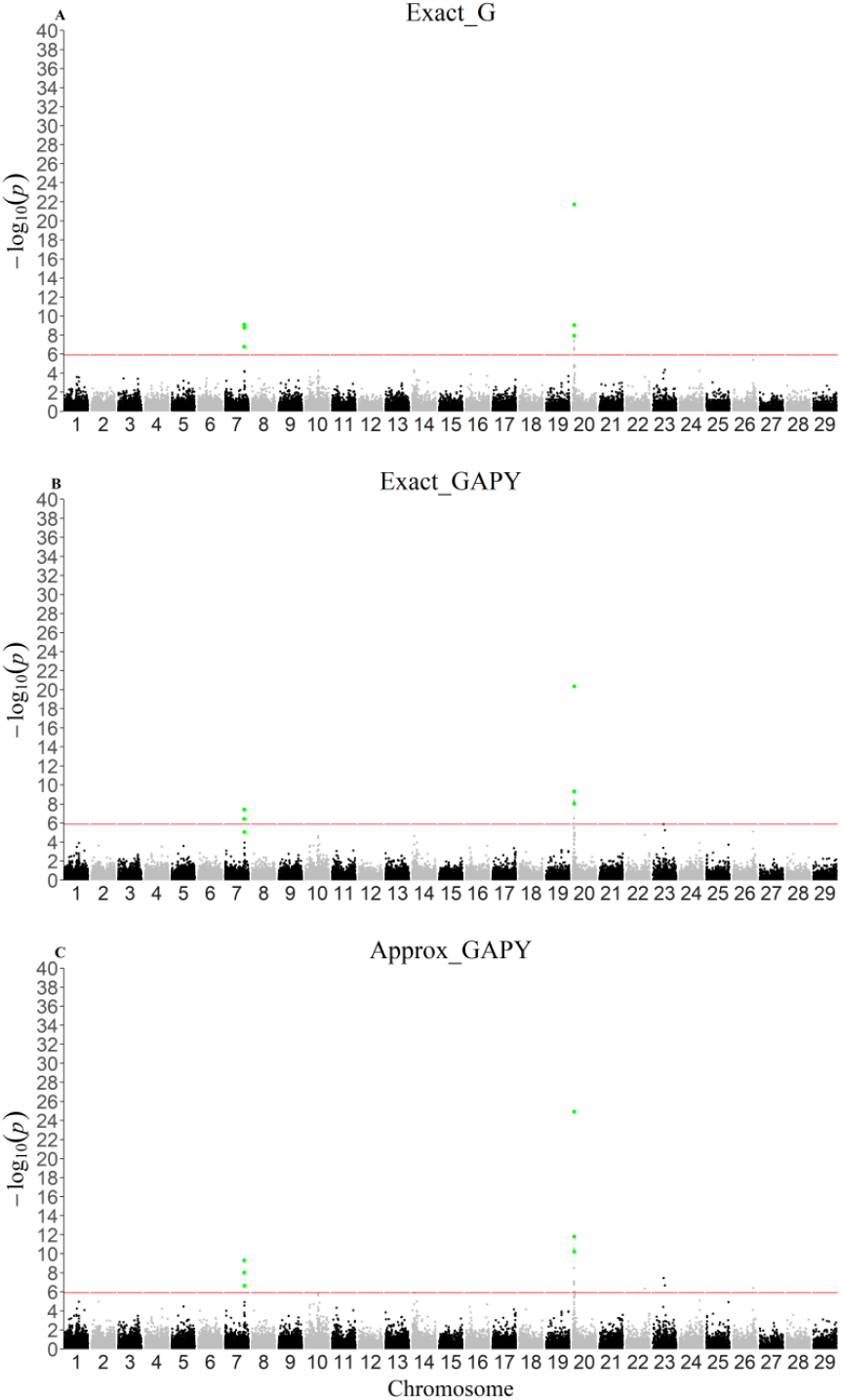
Manhattan plots for scenarios with a reduced data set in replicate 1. Single-step genome-wide association study for post-weaning weight with p-values obtained from a data set of 50K genotyped animals with (A) an exact inversion of the genomic relationship matrix (**G**^**−1**^; (Exact_Ginv), (B) a sparse **G**^**−1**^ build with the algorithm for proven and young (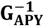; Exact_GinvAPY), and (C) an approximation based on 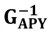 (Approx_GinvAPY) in replicate 1; SNPs highlighted in green represent the three most significant SNP in the two peaks found with Exact_Ginv.

**Figure 2.**
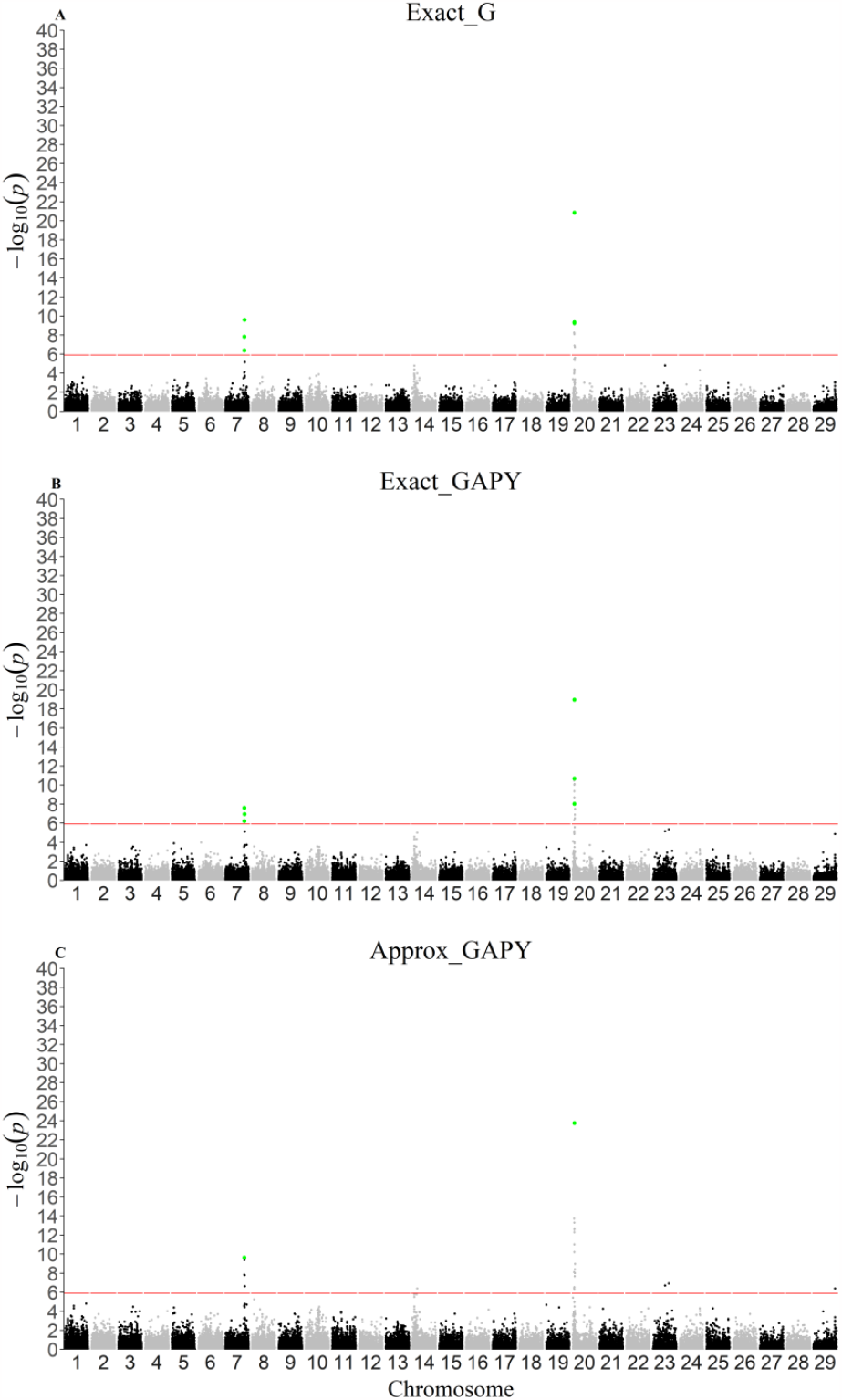
Manhattan plots for scenarios with a reduced data set in replicate 2. Single-step genome-wide association study for post-weaning weight with p-values obtained from a data set of 50K genotyped animals with (A) an exact inversion of the genomic relationship matrix (**G**^**−1**^; (Exact_Ginv), (B) a sparse **G**^**−1**^ build with the algorithm for proven and young (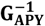; Exact_GinvAPY), and (C) an approximation based on 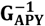 (Approx_GinvAPY) in replicate 2; SNPs highlighted in green represent the three most significant SNP in the two peaks found with Exact_Ginv.

**Figure 3.**
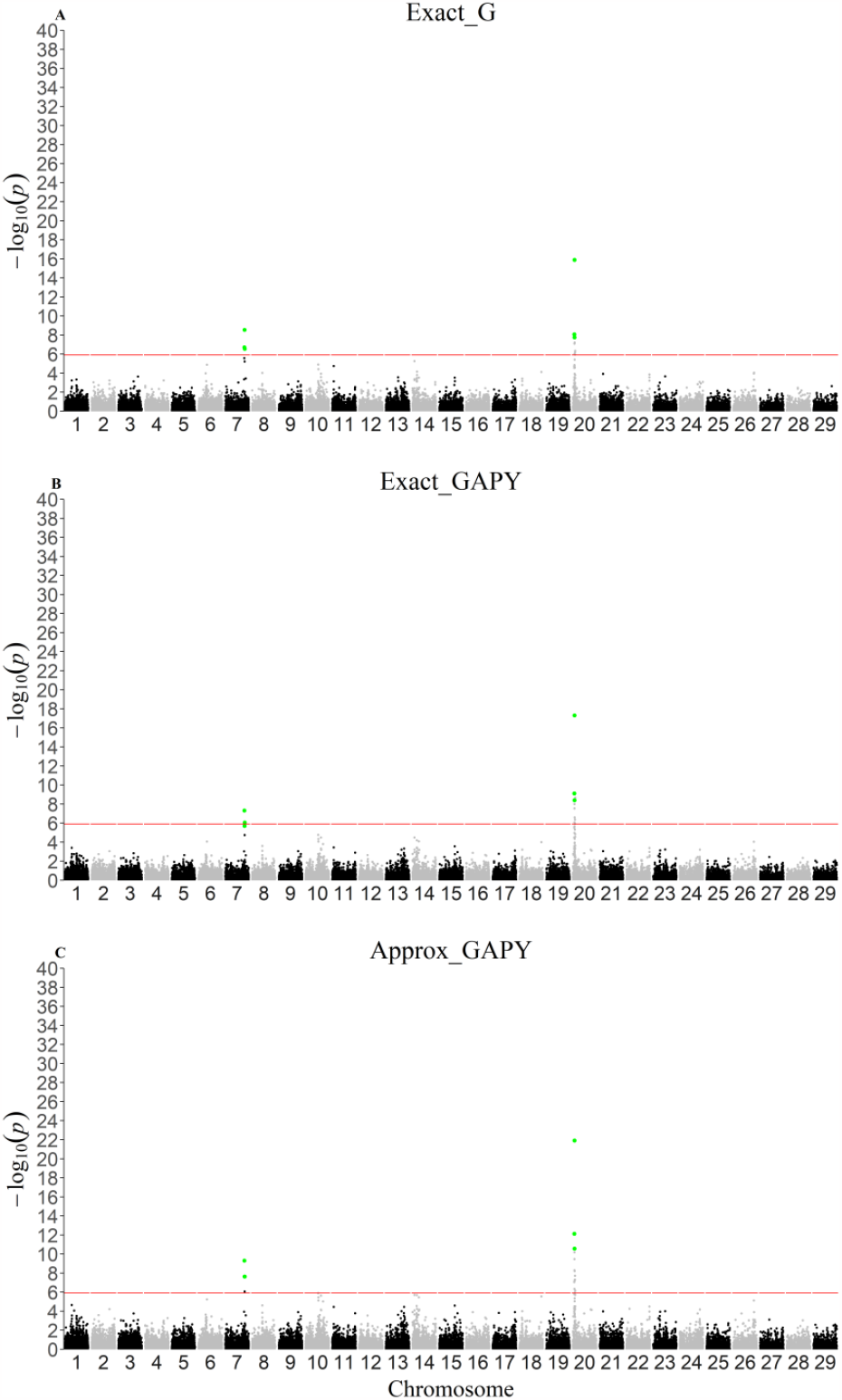
Manhattan plots for scenarios with a reduced data set in replicate 3. Single-step genome-wide association study for post-weaning weight with p-values obtained from a data set of 50K genotyped animals with (A) an exact inversion of the genomic relationship matrix (**G**^**−1**^; (Exact_Ginv), (B) a sparse **G**^**−1**^ build with the algorithm for proven and young (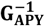; Exact_GinvAPY), and (C) an approximation based on 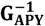 (Approx_GinvAPY) in replicate 3; SNPs highlighted in green represent the three most significant SNP in the two peaks found with Exact_Ginv.

P-values are functions of the SNP effect and their corresponding estimation variance (i.e., var(â_i_)). The correlation of those two variables between p-value computing methods is displayed in Figure 4. The correlation of SNP effects between Exact_Ginv and Approx_GinvAPY with respect to Exact_Ginv ranged from 0.87 to 0.89, whereas the correlation of var(â_i_) was slightly higher, ranging from 0.91 to 0.92, respectively (Figure 4). Between APY-based methods, the correlation for SNP effects and var(â_i_) approached unit (Figure 4). Although SNP effect correlations were overall high between all p-values computing methods, some SNPs in chromosomes 22, 23, 26, and 29 in Approx_GinvAPY achieved the significance threshold without a clear linkage disequilibrium trail, thus suggesting a slight increase in estimation noise in comparison to Exact_GinvAPY (Misztal et al., 2023). The increase in noise with Approx_GinvAPY may be explained by the two uncertainty measurements when obtaining p-values in this method. The first measurement of uncertainty is due to APY. The algorithm for proven and young is based on the theory that genomic information is limited, and that all genetic variation is contained in a set of independent chromosome segments within a population. Given that a core group of animals would contain those segments, the breeding values of noncore animals in the population could be estimated from the breeding values of core animals in addition to an error term 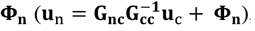, which is expected to approach zero when the number of SNP is equal or greater than the core size (Misztal, 2016; Bermann et al., 2022a). The second measurement of uncertainty comes from computing PEV_CCappx_. As shown by Misztal and Wiggans (1988), the off-diagonal elements are not considered during the absorption of environmental effects into the mixed model equations for constructing **D**. Thus, is expected that, with Approx_GinvAPY, there is a slight increase in noise, especially when the core set and data are small.

**Figure 4.**
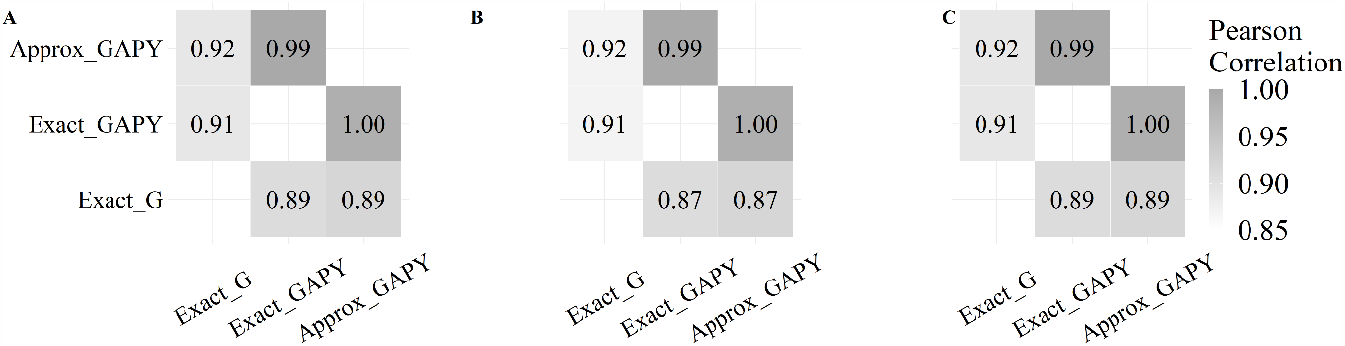
Person correlation between SNP effects and variance of estimated SNP effects across methods and replicates. Person correlation between SNP effects (upper-diagonal) and variance of estimated SNP effects (lower-diagonal) from different p-values computing methods in replicate 1 (A), replicate 2 (B), and replicate 3 (C). Methods refer to SNP p-values obtained from a data set of 50K genotyped animals with an exact inversion of the genomic relationship matrix (**G**^**−1**^; (Exact_Ginv), with a sparse **G**^**−1**^ build with the algorithm for proven and young (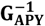; Exact_GinvAPY), and from an approximation based on 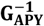 (Approx_GinvAPY). Approx_GinvAPY450K refers to the Approx_GinvAPY method applied to a genotyped population of 450K animals.

The elapsed wall-clock time and memory requirement across p-values computing methods are shown in Table 2, and despite ssGWAS results being similar among methods, computing time varied considerably. The average total elapsed wall-clock was 106.76 hours for Exact_Ginv, 110.98 hours for Exact_APY, and was reduced to 2.83 hours with Approx_GinvAPY (Table 2). Therefore, compared to Exact_Ginv, the run time with Approx_GinvAPY was reduced by about 38 times. The memory requirement also varied across p-values computing methods; its peak was 159.66 GB, 178.30 GB, and 16.62 GB for Exact_Ginv, Exact_APY, and Approx_GinvAPY, respectively. Compared to Exact_Ginv, the peak memory requirement for obtaining p-values with Approx_GinvAPY was about 10-fold smaller.

**Table 2.** Elapsed wall-clock time and memory requirement for p-values computation with Exact_Ginv, Exact_GinvAPY, and Approx_GinvAPY.

| Method<sup>a</sup> | Software | Elapsed time, h:min | SD | Peak of memory, GB<sup>b</sup> | SD |
| --- | --- | --- | --- | --- | --- |
| Exact_Ginv | **PREGSF90** | 1:01 | 0:27 | 107.28 | 0.00 |
|  | **BLUPF90IOD3** | 93.13 | 21:20 | 159.66 | 0.88 |
|  | **POSTGSF90** | 12:31 | 0:21 | 145.39 | 0.58 |
|  | **Total/Max** | 106:46 |  | 159.66 |  |
| Exact_GinvAPY | **PREGSF90** | 0:20 | 0:01 | 9.07 | 0.00 |
|  | **BLUPF90IOD3** | 108:45 | 22:33 | 178.30 | 0:00 |
|  | **POSTGSF90** | 1:53 | 0:39 | 44.83 | 0.00 |
|  | **Total/Max** | 110:59 |  | 178.30 |  |
| Approx_GinvAPY | **PREGSF90** | 0:39 | 0:16 | 9.07 | 0:00 |
|  | **BLUPF90IOD3** | 0:54 | 0:21 | 4.53 | 0:00 |
|  | **ACCF90GS2** | 0:04 | 0:02 | 9:07 | 0:00 |
|  | **POSTGSF90** | 1:50 | 0:06 | 16.62 | 0:00 |
|  | **Total/Max** | 2:50 |  | 16.62 |  |

<sup>a</sup>SNP p-values obtained from a data set of 50K genotyped animals with an exact inversion of the genomic relationship matrix ( $\mathbf{G}^{-1}$ ; (Exact\_Ginv), with a sparse $\mathbf{G}^{-1}$ build with the algorithm for proven and young ( $\mathbf{G}_{\text{APY}}^{-1}$ ; Exact\_GinvAPY), and from an approximation based on $\mathbf{G}_{\text{APY}}^{-1}$ (Approx\_GinvAPY).
<sup>b</sup>Resident Set Size (RSS) memory. Values are displayed as average and standard deviations among three replicates.

The most computationally demanding scenario was Exact_GinvAPY, with the computation of p-values taking a total wall-clock time run of 110.98 hours and a peak of memory requirement of 178.30 GB. Even though APY increases the sparsity of **G**^**−1**^ by ignoring the relationships between noncore animals, it still requires the storage of intermediate matrices and vectors. Moreover, the computational advantage with APY comes mainly from the block implementation with the preconditioned conjugate gradient (**PCG**) method, as shown by Masuda et al. (2016). However, in Exact_GinvAPY, the LHS is still explicitly inverted, which does not use the sparse properties of 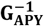 (Junqueira et al., 2022). Although Exact_GinvAPY has a similar computing performance as Exact_Ginv, results from this method are helpful in this study to illustrate the feasibility of accurately computing p-values with 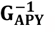.

### Application of Approx_GinvAPY with a large genotype dataset

Manhattan plots for p-values obtained with Approx_GinvAPY450K are in Figure 5. Across all replicates, the two significant peaks in chromosomes 7 and 20 observed in the first set of analyses with Exact_Ginv (benchmark) were also identified with Approx_GinvAPY450K. Apart from those two persistent peaks, two new peaks in chromosomes 6 and 14 were uncovered with Approx_GinvAPY450K (Figure 5). The new peaks had clear linkage disequilibrium trails, meaning that ssGWAS resolution increases as more genotyped animals are included in the analyses. As previously shown, especially for populations with a small effective population size (**Ne**) and more polygenic traits, increasing the genotype set reduces the estimation error and the shrinkage of SNP effects, which increases the power of discovering significant variants (Lourenco et al., 2017; Jang et al., 2023; Misztal et al., 2023). The benefit of an increase in the genotype set size can also be observed when comparing Approx_GinvAPY with approx_GinvAPY450K. In general, for significant SNPs identified on chromosomes 7 and 20, the magnitude of p-values on the logarithmic scale obtained with Approx_GinvAPY450K increased by 50% relative to results from Approx_GinvAPY.

**Figure 5.**
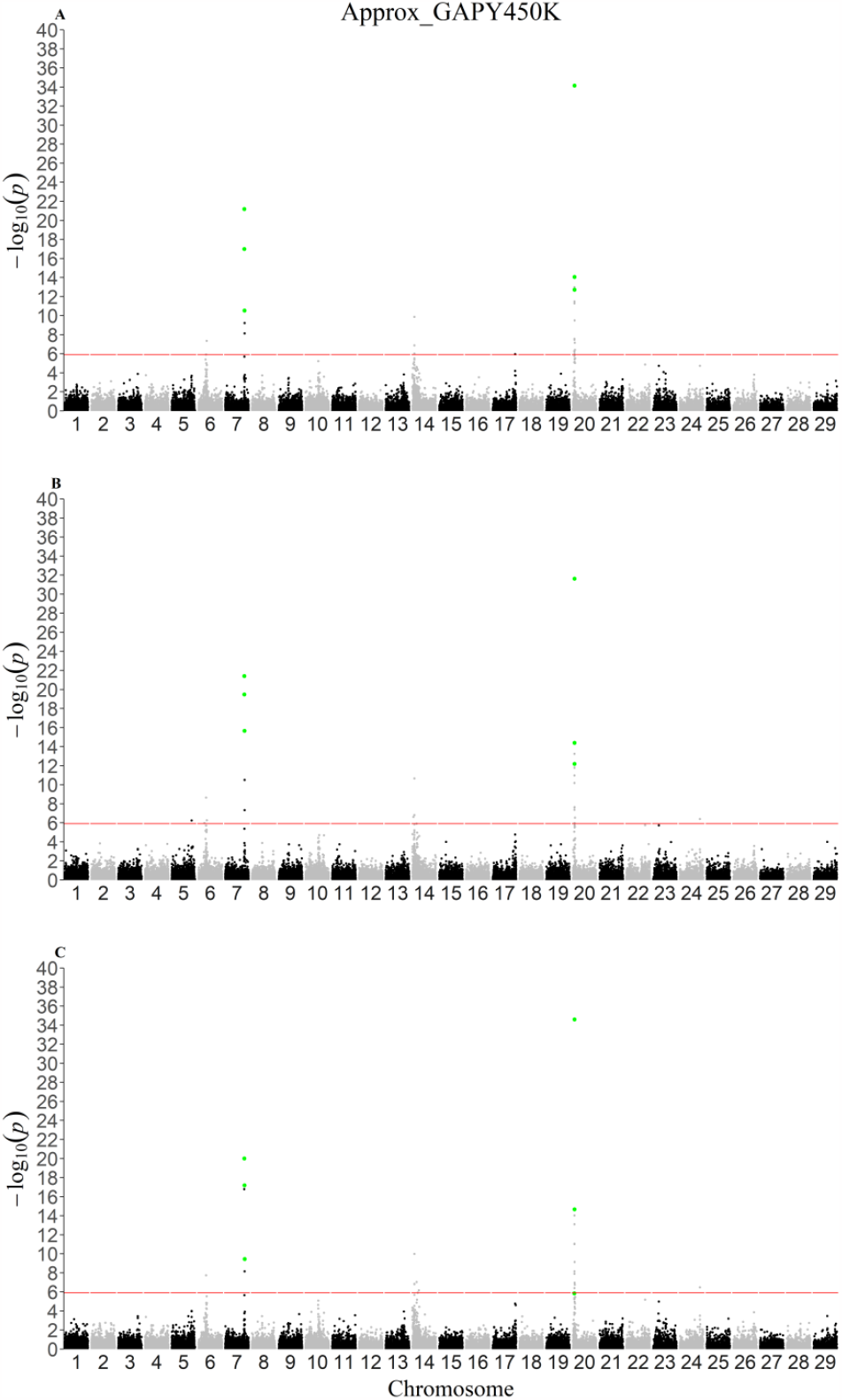
Single-step genome-wide association study for post-weaning weight using Approx_GinvAPY450K across all replicates. Single-step genome-wide association study for post-weaning weight using Approx_GinvAPY450K in (A) replicate 1, (B) replicate 2, and (C) replicate 3; SNP highlighted in green represent the three most significant SNP in the two peaks found in with Exact_Ginv with a reduced genotype dataset. Approx_GinvAPY450K refers to the method where SNP p-values obtained from a data set of 450K genotyped animals with an approximation based on 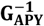.

In the first set of analyses, when the same amount of data was used, an increase in noise was observed with Approx_GinvAPY compared to Exact_GinvAPY (Figures 1-3). However, when more genotyped animals were included with Approx_GinvAPY450K, significant SNPs without a clear LD pattern were no longer observed in all replicates (Figure 5). This suggests that the benefit of increasing the genotype set overcomes the noise associated with an approximation that relies on APY and PEV_CCappx_ and mitigates potential false positive associations. While evaluating two simulated populations with the same Ne, Misztal et al. (2023) observed that increasing the number of individuals contributing with genotypes and phenotypes by three times increased the correct identification of significant SNPs. Similarly, Jang et al. (2023) showed that for highly polygenic traits (2000 QTN) with Ne of 20 and a moderate heritability of 0.30, no QTN was accurately identified until a complete genotype set, composed of 30K genotyped animals, was included in the analyses. For livestock populations with even smaller Ne and traits of lower heritability, such as reproduction and fitness traits, QTN identification may be even more challenging, especially when limitations exist on the amount of genomic information used in the estimation process.

Total wall-clock time for the calculation of p-values with Approx_GinvAPY450K was, on average, 24.47 hours, which was divided into building 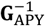 and saving components of 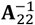 (6.6 hours), estimation of breeding values (6.67 hours), estimation of PEV_CCappx_ (0.38 hours), and backsolving GEBV to SNP effects and approximation of var(â_i_) (Table 3). The entire process required no more than 87.64 GB of memory (Table 3). In comparison with the same method using a reduced set of genotyped animals in the first set of analyses (i.e., Approx_GinvAPY), the increase in wall-clock time was linear with the increase in the number of genotypes included added, which was approximately nine times. However, the increase in memory requirement was only five times.

**Table 3.**
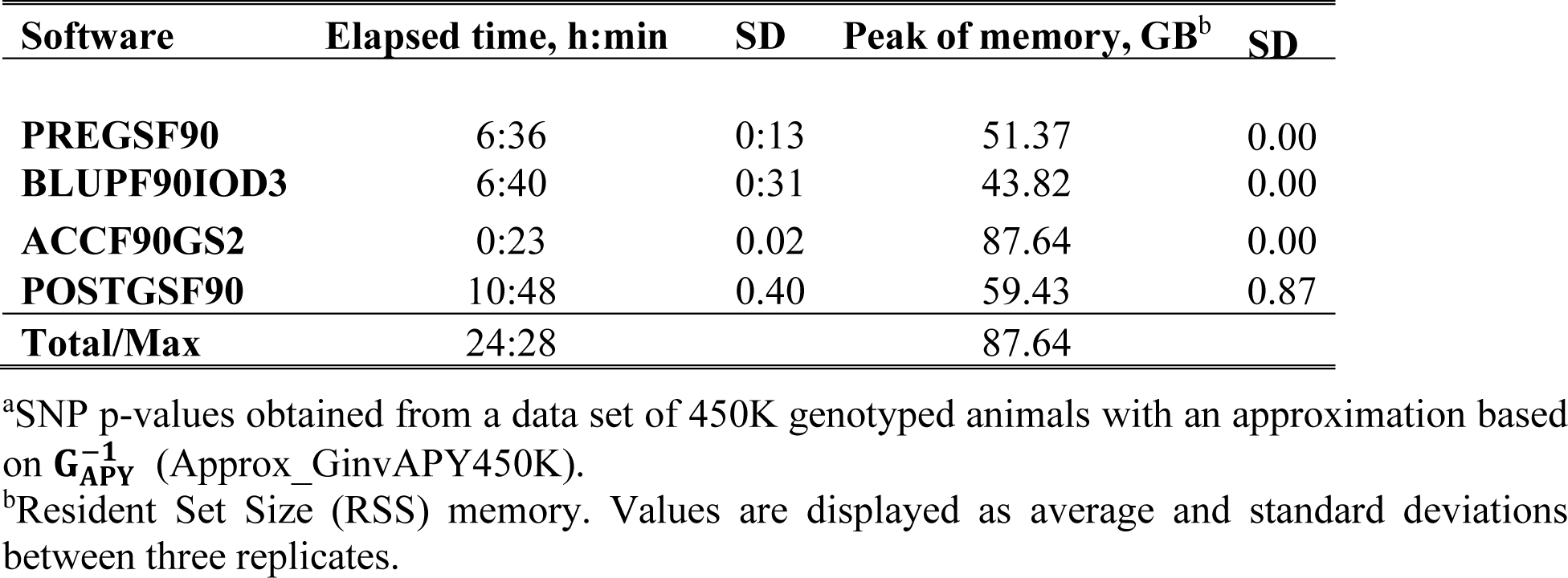
Elapsed wall-clock time and memory requirement for p-values computation with Approx_GinvAPY450K^a^.

The efficiency of the proposed approximation method (i.e., Approx_GinvAPY) is because var(â_i_) computations rely only on the genotypes of core animals, meaning that the computational requirement of inverting **G** in Exact_Ginv and obtaining 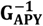 in Exact_GinvAPY is reduced to inverting a small matrix of relationships between core animals (**G**_**cc**_) (Bermann et al., 2022a). The dimension of **G**_**cc**_ is approximately a linear function of Ne of the population and should not be more than 15K for most livestock species or breeds (Pocrnic et al., 2016; Cesarani et al., 2022). Moreover, with the Approx_GinvAPY method, no inversion of the LHS is required. Instead, PEV_CCappx_ are obtained accurately with a lower computational cost by a block sparse inversion of 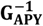that had weights (effective record contributions; **D** in Eq. [9]) added to its diagonal (Bermann et al., 2022b). Additionally, because Approx_GinvAPY does not require the direct inversion of the LHS, efficient solvers such as PCG can be used in combination with the block implementation of APY, efficiently exploiting the sparseness of **G**^**−1**^ (Masuda et al., 2016; Junqueira et al., 2022).

Altogether, our results show that the current computational limitations for obtaining p-values for populations with many genotyped animals are no longer an issue with Approx_GinvAPY. The possibility of computing SNP p-values for those large genotyped populations should increase the power of detection of true variants and prevent future findings of ssGWAS from solely relying on SNP frequency and effect in the population (Wang et al., 2012; Aguilar et al., 2019). However, it is worth noting that the results presented herein are based on a single-trait model in a purebred population. As long as reliabilities from more complex models and on populations with more complex breeding structures are accurately estimated, we expect that p-values will also be accurately approximated. However, such scenarios deserve further investigation.

## Conclusions

The same genomic regions in chromosomes 7 and 20 were identified with p-values obtained with **G**^**−1**^, 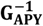, and the approximation based on 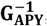 with a reduced dataset, indicating the soundness of the proposed algorithm. Even though p-values were similar between computing methods, computational requirements for the new algorithm were considerably reduced. When the approximation based on 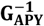 was applied to a genotyped population with almost half a million genotyped animals, SNPs on chromosomes 7 and 20 had stronger signals, and two new regions on chromosomes 6 and 14 were uncovered, indicating an increase in ssGWAS detection power when more genotypes are included in the analyses. Obtaining p-values in ssGWAS for such a large genotyped population required 24 hours, which is expected to increase linearly with the addition of noncore genotyped individuals. With a combination of APY and an approximation of the variance of estimating SNP effects, ssGWAS with p-values becomes computationally feasible for large genotyped populations.

## Acknowledgments

We thank the American Angus Association (St. Joseph, MO) for conceding the dataset used in this study.

